# SentryPath: a mechanistic protocol-ranking simulator with leave-one-trial-out cross-validation across 13 phase-III oncology randomised controlled trials and a pre-registered prospective forecast

**DOI:** 10.64898/2026.06.23.734048

**Authors:** M Deeraj Kumar, S Manoj Kumar

## Abstract

**Background:** Pivotal oncology trials cost a median of ≈$19 million each (oncology often $45 million or more) and contribute to a capitalised cost of ≈$2.6 billion per approved drug, yet most candidate protocols never reach trial. Existing in-silico screening tools either rely on closed proprietary PK/PD modelling or require patient-level data; a transparent, cohort-level, cross-validated mechanistic alternative is missing.

**Methods:** SentryPath is a physics-based stochastic differential equation simulator built on a Gompertzian tumour-growth term with Emax pharmacodynamic kill and Bliss-independence combination modelling, scored at the cohort level. Validation against 13 published phase-3 randomised controlled trials covering six cancer types uses the 2-year overall-survival (OS) rate ratio as the primary endpoint, cross-checked against ClinicalTrials.gov posted results. For cancer types with ≥2 trials we apply leave-one-trial-out crossvalidation: two shared efficacy scalars per cancer type are fit on training trials and used to predict the heldout trial cold.

**Results:** With the per-drug efficacy proxies held fixed from the literature, two shared cancer-type scalars fit on the training trials transfer to the held-out trial with a **mean held-out error of 3.7 % (range 0.7–7.3 %)** on 2-year OS rate ratios across three NSCLC trials; extending the same method to RCC, HCC, and ESCC yields a 5.4 % aggregate across nine folds (per-fold range 0.2–11.2 %), reported with per-cancer stratification. We are explicit that only the two scalars are held out — the per-protocol efficacy proxies underneath are literature-anchored to drug classes that include the benchmark trials, so this is a test of scalar transfer, not of the whole engine cold. Cross-validation improves on the same engine without it (16.4 % with production cancer priors; 21.9 % with no efficacy modifiers); a matched in-sample fit of the same two-scalar model gives 4.4 %, slightly below the 5.4 % held-out, the expected direction. Two prospective forecasts are preregistered on the Open Science Framework with falsification envelopes and pre-readout bias disclosure. The first forecast (NCT04770896) reaches its primary data cutoff on 2026-06-30; the observed outcome and its mapping to the pre-committed interpretation will be reported in a versioned update to this preprint.

**Conclusion:** A transparent mechanistic simulator, with a literature-anchored efficacy library and only two cross-validated scalars per cancer type, transfers those scalars across held-out NSCLC trials at 3.7 % mean error (range 0.7–7.3 %) and extends to other cancers with documented per-cancer stratification. The validation is pilot-scale (3–9 folds) and the scalars sit on a fixed, trial-informed substrate; its distinguishing contribution is less the error magnitude than the public predict–verify–disclose cycle that goes beyond retrospective fit.

## 1. Introduction

### 1.1 The problem

Pivotal phase-3 oncology trials are among the most expensive experiments in medicine. Across 138 pivotal trials supporting US FDA approvals in 2015–2016, the median estimated trial cost was $19 million (interquartile range $12–33 million), with oncology trials among the most expensive (often exceeding $45 million) and a more-than-100-fold spread overall (Moore et al. 2018). Timelines run three to five years from launch to primary readout. These per-trial figures are a small fraction of the capitalised cost per approved drug — estimated at ≈ $2.6 billion including the cost of failures and of capital (DiMasi et al. 2016) — and the dominant driver of that total is attrition (Wong et al. 2019). The attrition is severe: the large majority of oncology agents that enter clinical testing never reach approval, and a substantial share of late-stage failures involve a protocol decision — a combination, a dose schedule, a line of therapy, a population — that did not deliver the relative benefit its sponsors expected. Many such failures are, in principle, knowable before the trial is run. What is missing is a cheap, transparent, and trustworthy way to screen candidate protocols *before* the phase-3 commitment, at the level of cohort-scale relative outcomes rather than individual patients.

### 1.2 Existing approaches and their gaps

Three families of in-silico tools address adjacent problems, but none fill this specific gap.

#### Regulatory-grade PK/PD platforms

Physiologically-based pharmacokinetic and population PK/PD modelling (e.g. Certara/Simcyp) is the established standard for predicting drug exposure and is widely accepted by regulators. These platforms are built for pharmacokinetics, however, not for ranking protocol-level *outcomes* such as the survival benefit of one regimen over another, and they are closed-source and costly.

#### Machine-learning digital-twin frameworks

Patient-level approaches — prognostic digital twins (Walsh et al. 2020) and the broader data platforms typified by Tempus and Foundation Medicine — can be powerful where rich patient-level data exist. But they require exactly that data, are difficult to inspect or audit, and cannot be deployed for a candidate protocol whose population has not yet been enrolled.

#### Bespoke mechanistic oncology modelling

Mechanistic tumour-dynamics models in the tradition of Gompertzian growth and Emax pharmacodynamics (Laird 1964; Benzekry et al. 2014; Enderling & Chaplain 2014) are biologically interpretable and need no patient-level data. In practice, however, they are typically delivered as per-project consulting, are rarely validated across more than one cancer type, and rarely commit to predictions in advance of the data they aim to anticipate.

### 1.3 What is missing

The unoccupied position is a single tool that is simultaneously transparent and open, cohort-level (requiring no patient-level data), parsimonious, validated by cross-validation across multiple cancer types, and prospectively pre-registered. SentryPath is built to occupy that position. It is a physics-based stochastic differential equation simulator of tumour-burden trajectory under therapy, combining Gompertzian growth, an Emax/Hill pharmacodynamic kill term, Bliss-independence combination modelling (Bliss 1939; Berenbaum 1989), and immune, resistance, and toxicity dynamics, scored at the cohort level against published 2-year overall-survival rate ratios. We emphasise at the outset that SentryPath *simulates, estimates, and ranks* protocols for exploratory screening; it is not a digital twin and not a clinical decision aid, and the validation that follows concerns relative outcomes between trial arms, not absolute survival or individual-patient prediction.

### 1.4 Contributions

This paper makes three specific contributions, each delivered by a later section.

1. **A multi-cancer retrospective validation with independent verification**. We validate the engine against 13 published phase-3 randomised controlled trials spanning six cancer types, using the 2-year overall-survival rate ratio between experimental and control arms as the primary endpoint, with every target ratio cross-checked against ClinicalTrials.gov posted-results data (§2.5, §3.3–§3.4).
2. **A leave-one-trial-out cross-validation framework for the cancer-type scalars**. With the per-protocol efficacy proxies held fixed, two shared efficacy scalars per cancer type — fit on training trials and applied cold to a held-out trial — reproduce held-out 2-year OS rate ratios with a mean error of **3.7 % (range 0.7–7.3 %) across the three NSCLC trials**, and 5.4 % aggregated across nine folds and four cancer types, reported with explicit per-cancer stratification (§2.6, §3.1–§3.2). The cross-validated scalars improve on cancer-type production priors used without cross-validation (16.4 %; §3.7). We are explicit (§3.7, Supplementary Table S1) that only the two scalars are cross-validated — the literature-anchored efficacy substrate beneath them is not held out — so the claim is scalar transfer across trials, not the whole engine generalising cold.
3. **A prospective pre-registration with public bias-disclosure**. We lock prospective trial forecasts on the Open Science Framework — timestamped, hash-fingerprinted, and falsifiable — together with a pre-readout disclosure of expected directional bias and a pre-committed mapping from each possible outcome to a fixed methodological response (§2.8, §3.6, §4.3). The first such forecast resolves on 2026-06-30.

Together these move the contribution beyond retrospective fit toward a transparent, parsimonious, crossvalidated methodology that commits to its predictions before the evidence arrives.

## 2. Methods

### 2.1 Engine architecture

SentryPath is a stochastic differential equation (SDE) simulator of tumour-burden trajectory under therapy. The full pipeline — from the protocol library and synthetic cohort, through the mechanistic ODE core, to per-protocol statistics and the ranked output — is shown in Figure 1.

**Figure 1:**
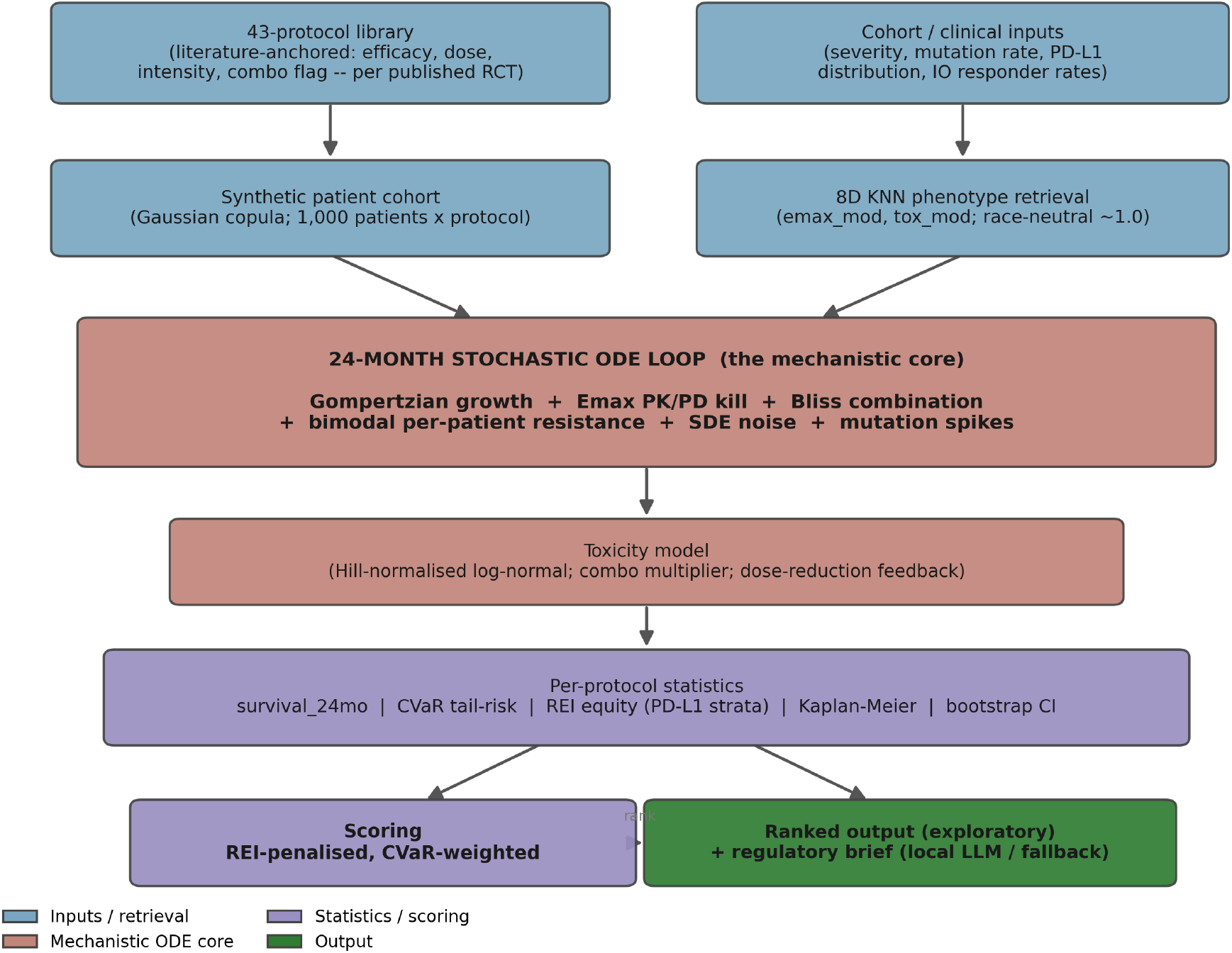
SentryPath simulation pipeline (population mode). Each stage is a deterministic, seed-locked transform; no neural network.

For each of *N* synthetic patients in a cohort (default *N* = 1,000 per protocol, scaling to 43,000 across the 43-protocol library; Section 2.4), the engine integrates a Gompertzian growth term coupled to an Emax (maximum-effect) pharmacodynamic kill term, perturbed by a Wiener process to capture biological variability:

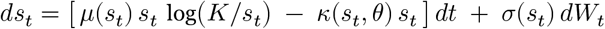

where *s*_*t*_ ∈ [0, 1] is the normalised tumour burden at month *t, K* = 1.01 is a carrying-capacity proxy, *μ*(⋅) is the intrinsic growth coefficient (a function of patient severity and mutation rate), *⋅*(⋅, *θ*) is the protocol-dependent kill rate, *σ*(⋅) is the patient-specific biological-variability term, and *W*_*t*_ is standard Brownian motion. The simulation is integrated explicitly over *T* = 24 discrete monthly time-steps using d*t* = 1 month.

#### 2.1.1 Growth term

The intrinsic Gompertzian coefficient *μ* for each patient is derived from cohort-level severity *S* ∈ [0, 1] and population mutation rate *M* ∈ [0, 1]:

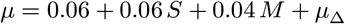

where *μ*_Δ_ is an optional hereditary modifier (BRCA, TP53, Lynch syndrome, etc.) applied in single-patient mode. The Gompertzian form (Laird 1964; Benzekry et al. 2014) is well-validated for solid tumours and naturally captures the slowing growth as *s*_*t*_ → *K*.

#### 2.1.2 Pharmacodynamic kill term (Emax/Hill)

The protocol-dependent kill rate uses the Hill equation:

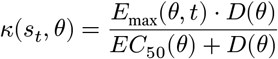

where *D*(*θ*) is the protocol’s normalised dose intensity, *EC*_50_ is the dose for half-maximal effect, and the time-varying *E*_max_ aggregates a stack of biologically interpretable multipliers:

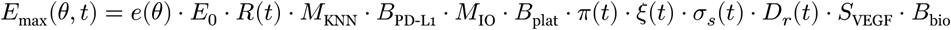

with terms defined as: - *e*(*θ*) is the protocol’s literature-anchored efficacy proxy (∈ [0.28, 0.90], Section 2.4); - *E*_0_ = 0.12 is `EMAX_SCALE`, the single global calibration constant fit on CheckMate 057 (Borghaei et al. 2015); - *R*(*t*) is the resistance decay term modelling acquired resistance after month 12; - *M*_KNN_ is a perpatient pharmacogenomic modifier from a *k*-nearest-neighbour retrieval against a synthetic twin database (race-neutral; see §2.2); - *B*_PD-L1_ is a boost dependent on programmed death-ligand 1 (PD-L1) expression stratum for immuno-oncology (IO)-class protocols (1 + 0.20 ⋅ PD-L1/2); - *M*_IO_ is an IO-class multiplier for hereditary modifiers (e.g. Lynch syndrome MSI-high ×1.45); - *B*_plat_ is a platinum-class booster for BRCA carriers (+0.30 to +0.45 Emax); - *π*(*t*) = exp(−*k*_*e*_ ⋅ *t*) is PK clearance with drug-class-specific *k*_*e*_ ∈ {0.005, 0.012, 0.020} per month; - *ξ*(*t*) is an immune-exhaustion factor activating after 12 months of sustained IO response (1.8 % per-month decay, capped at 45 % reduction); - *σ*_*s*_(*t*) is a protocol schedule multiplier capturing induction–maintenance dynamics; - *D*_*r*_(*t*) = 1 − 0.30 ⋅ max(0, (*τ* (*t*) − 0.70)/0.30) is a toxicity-feedback dose-reduction factor (Grade 3/4 patients receive up to 30 % Emax reduction); - *S*_VEGF_ is an IO + anti-VEGF synergy boost (+0.15, gated against double-counting in benchmarks already calibrated for this synergy); - *B*_bio_ is a biomarker resistance-gene modifier (STK11 ×0.80, KEAP1 ×0.85, MSI-H ×1.35 for IO arms, clamped at 0.60 floor).

The stacked-multiplier form is biologically motivated: each multiplier corresponds to a published mechanism (cited in Supplementary Table S1), and the floor-clamping prevents pathological compounding.

#### 2.1.3 Combination therapy (Bliss independence)

For combination regimens (`combo = 1` in the protocol library), the kill term is computed via Bliss independence on fractional effects:

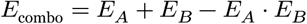

where *E*_*A*_, *E*_*B*_ ∈ [0, 1] are the fractional kill effects of each component arm. This is the standard pharmacological assumption when component mechanisms are independent (Bliss 1939; Berenbaum 1989). The model’s known underestimation of chemo-IO synergy (Galluzzi 2020) is documented as a limitation (§4.4) with a planned upgrade discussed in §4.7.

#### 2.1.4 Stochastic perturbation

The SDE noise term *σ*(*s*_*t*_) = 0.05 + 0.05 ⋅ *s*_*t*_ scales with burden, capturing greater variability in higher-burden patients. An additional Bernoulli failure spike (`failure_prob` ∈ [0.001, 0.04] depending on severity and intensity) adds a +0.10 burden shock to model rare mutation-driven progression events.

#### 2.1.5 Lethality threshold and competing-risk mortality

Time-to-event survival is derived from the lethality threshold *s*_*t*_ ≥ 0.80. Patients who do not cross this threshold within 24 months are censored at study end. A competing-risk background mortality of ∼ 3.6% per year (exponential, scale 333 months) is added independently to capture non-cancer-attributable deaths in the 60–70-year-old oncology population (matching the published ∼ 7–15% non-cancer trial-mortality range).

### 2.2 Patient cohort generation

Each simulation run generates a synthetic cohort of 1,000 patients per protocol via independent sampling from a parameterised distribution. The cohort is fully characterised by an 8-dimensional patient feature vector and a small set of cohort-level priors.

#### 2.2.1 Eight-dimensional patient vector

For each patient *i* ∈ {1, …, *N*}:

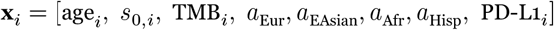

- Age ∼ *N*(60, 10) years
- Initial burden *s*_0,*i*_ ∼ *N*(*S, τ*) clamped to [0.05, 0.95], with *S* = cohort severity and *τ* = cohort burden standard deviation
- Tumour mutational burden (TMB) ∼ *N*(15, 5) mutations/Mb
- Ancestry one-hot encoded across {European, East Asian, African, Hispanic}, flat 25 % distribution
- PD-L1 stratum drawn from cohort-specific distribution [neg, low, high] summing to 1

#### 2.2.2 Race-neutral pharmacogenomic retrieval (v4.2 onwards)

Earlier versions of SentryPath applied ancestry-indexed pharmacokinetic multipliers (European 0.95 / East Asian 1.15 / African 0.85 / Hispanic 1.05 on Emax) ostensibly motivated by CYP2C19/CYP3A5 allelefrequency distributions. **These race-based scalars were removed in v4.2** because they encoded an algorithmic-harm pattern (race → drug efficacy) that contradicts the project’s own equity-aware design (Obermeyer et al. 2019). The ancestry features remain in the patient vector for future genotype-grounded use, but the KNN-retrieved Emax modifier *M*_KNN_ is now uniform across ancestry (1.0 + Gaussian noise). Race is not an input to efficacy.

Per-patient pharmacogenomics, when relevant, is now treated as an explicit per-patient genotype input (CYP2C19 *2/*3, CYP3A5 *1, BRCA, TP53, Lynch, etc.) rather than a population-race proxy.

#### 2.2.3 Cohort priors and biomarker layers

Each benchmark cohort is characterised by: - Severity *S* (derived from published median OS of the control arm in the source trial) - Mutation rate *M* (mapped from published TMB category) - PD-L1 distribution (mapped from published PD-L1 enrichment in the trial population) - IO complete-response rate (*io*_CR_) and durable-response rate (*io*_DR_): population-level Bernoulli sampling assigning each IO-arm patient to complete responder / durable responder / progressor categories with literature-anchored fractions per cancer type - Resistance-gene rates (STK11, KEAP1, MSI-H): per-patient Bernoulli draws using cohort-specific prevalence rates

Full per-cohort priors are listed in Supplementary Table S2.

### 2.3 Scoring framework

A protocol’s composite score is:

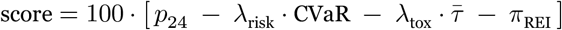

where *p*_24_ is the cohort-mean 24-month survival probability, the conditional value-at-risk (CVaR) captures tail risk, 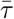 is mean toxicity, *π*_REI_ is the representation equity index (REI) biomarker subgroup-equity penalty, *λ*_risk_ is the user-set risk-aversion coefficient (∈ [0, 1]), and *λ*_tox_ = 0.3 is fixed.

#### 2.3.1 CVaR tail-risk

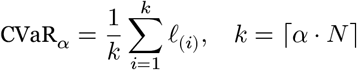

where *ℓ*_(*i*)_ is the *i*-th largest patient loss (defined as (*T* − *t*_event,*i*_)/*T* for died patients, 0 for censored). The risk-aversion-modulated *α* = 0.20 − 0.15 ⋅ *λ*_risk_ (conservative *λ*_risk_ = 1 gives the worst-5 % tail). CVaR penalises protocols that produce concentrated early mortality, even if their mean survival is acceptable.

#### 2.3.2 Representation Equity Index (REI)

REI measures treatment-effect heterogeneity across PD-L1 biomarker strata (Negative / Low 1–49 % / High ≥50 %):

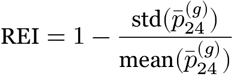

across PD-L1 strata *g* present in the cohort. A protocol whose benefit concentrates in PD-L1-high patients (leaving PD-L1-negative behind) scores a lower REI than one with even cross-stratum benefit. The equity penalty is:

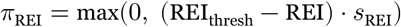

with REI_thresh_ = 0.95 (the 25th percentile of the empirical REI distribution across 120 protocol × cohort observations) and *s*_REI_ = 3.0 (rescaling for the compressed metric range). The threshold and scale were recalibrated in v4.2 after the metric was re-defined from ancestry-based to biomarker-based grouping.

#### 2.3.3 Pareto-optimality filter

Independently of the composite score, protocols are tagged as Pareto-optimal in the (survival, toxicity) plane: a protocol is Pareto-optimal if no other evaluated protocol simultaneously achieves higher mean survival AND lower mean toxicity. Pareto-optimal protocols are surfaced alongside the score-ranked list.

#### 2.3.4 Hard contraindication gating

Independently of scoring, a clinical contraindication module (v4.2) removes protocols a patient cannot safely receive from the ranked recommendation set: - Child-Pugh B/C → exclude bevacizumab-containing regimens (variceal-bleeding risk; IMbrave150 enrolled CP-A only) - Active autoimmune disease → exclude checkpoint-blockade-containing regimens (severe irAE flare risk; NCCN guidance) - Creatinine clearance < 60 mL/min → exclude cisplatin-containing regimens (nephrotoxicity) - ECOG ≥ 3 → exclude intensive cytotoxic/combination regimens

Each exclusion is logged with a human-readable reason; the contraindicated set is returned alongside the eligible recommendations.

### 2.4 Protocol library

The simulator includes 43 literature-anchored regimens spanning the major drug classes used in 1L and 2L oncology trials across the six covered cancer types. Each protocol is specified by four parameters (efficacy, dose intensity, kinetics category, and combination flag), all derived from published phase II/III RCT endpoints with full trial citation — and several of those source trials are also benchmark trials, so the base efficacy proxies are not blind to the benchmark suite. The base protocol library is fixed at simulation start and identical across all benchmark runs; the per-trial *class multipliers* applied on top of it (and their full degrees-of-freedom audit) are reported in Supplementary Table S1, and the cross-validated component is restricted to the two cancer-type scalars of §2.6.

Full protocol-by-protocol parameters and source citations are listed in Supplementary Table S1.

### 2.5 Validation framework

#### 2.5.1 Benchmark suite (13 phase-III RCTs)

| # | Trial | Cancer | NCT | Primary citation |
| --- | --- | --- | --- | --- |
| 1 | CheckMate 057 (CALIBRATION) | NSCLC 2L | NCT01673867 | Borghaei NEJM 2015 |
| 2 | KEYNOTE-024 | NSCLC 1L<br>PD-L1 ≥ 50% | NCT02142738 | Reck NEJM 2016 / JCO 2021 |
| 3 | KEYNOTE-189 | NSCLC 1L all<br>PD-L1 | NCT02578680 | Gandhi NEJM 2018 |
| 4 | CheckMate 214 | RCC 1L int/poor | NCT02231749 | Motzer NEJM 2018 |
| 5 | KEYNOTE-426 | RCC 1L all-risk | NCT02853331 | Rini NEJM 2019 / Lancet Onc 2020 |
| 6 | CheckMate 066 | Melanoma 1L | NCT01721772 | Robert NEJM 2015 |
| 7 | CheckMate 067 | Melanoma 1L | NCT01844505 | Larkin NEJM 2019 / Wolchok NEJM 2025 |
| 8 | IMbrave150 | HCC 1L | NCT03434379 | Finn NEJM 2020 |
| 9 | HIMALAYA | HCC 1L | NCT03298451 | Abou-Alfa NEJM Evid 2022 |
| 10 | KEYNOTE-590 | ESCC 1L | NCT03189719 | Sun Lancet 2021 |
| 11 | CheckMate 648 | ESCC 1L | NCT03143153 | Doki NEJM 2022 |
| 12 | TOPAZ-1 | BTC 1L | NCT03875235 | Oh NEJM Evid 2022 |
| 13 | KEYNOTE-966 | BTC 1L | NCT04003636 | Kelley Lancet 2023 |

CheckMate 057 is the sole calibration trial — `EMAX_SCALE = 0.12` and the immune-exhaustion decay (1.8 %/month) were jointly fit on its published 2-year OS rate ratio (Nivolumab/Docetaxel = 0.29/0.16 = 1.813). The remaining 12 trials are out-of-calibration-distribution.

#### 2.5.2 Primary endpoint: 2-year OS rate ratio

For each trial, the primary validation metric is the ratio of 2-year overall-survival rates between the experimental and control arms:

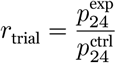

The 2-year landmark was chosen because (a) it is the most commonly reported survival landmark across our 13 trials, (b) it sits within the simulator’s 24-month time-step window without extrapolation, and (c) ratios are scale-invariant and less sensitive to absolute-survival calibration than hazard ratios (the simulator’s HR is structurally compressed; see §4.4). Where the source publication does not report the 24-month landmark directly, the per-arm rate is derived from published median OS via an exponential-survival model (§2.5.3); this derivation is exact only for exponential survival and is least accurate for plateauing immunotherapy curves, a limitation carried forward to §4.4.

#### 2.5.3 Target ratio derivation and verification audit

For each trial we extract the published 2-year OS rate per arm from the source publication or, where the source does not report a 24-month landmark explicitly, derive it from the published median OS via the exponential model *S*(24) = exp(− ln 2 ⋅ 24/mOS) (consistent with network-meta-analysis methodology).

On 2026-05-31 (v4.3.1) we systematically cross-checked every target ratio against the ClinicalTrials.gov v2 REST API posted-results data for the corresponding NCT identifier. The audit identified one materially-biased target (HIMALAYA: 1.33 → 1.242; the original value had been derived from 3-year OS data rather than the 24-month landmark) and two minor refinements (CheckMate 066: 2.161 → 2.198; CheckMate 067: 1.10 → 1.085).

Five trials (CheckMate 057, 066, 067, 214, HIMALAYA) are now directly verified against CT.gov posted-results data (Tier-A). The remaining eight trials have published median OS and HR confirmed against CT.gov but require primary-publication figure-reading of the 24-month Kaplan–Meier landmark to be fully verified (Tier-B); per-trial verification status is reported in Supplementary Table S3.

#### 2.5.4 Error metric

Per-trial error is computed as

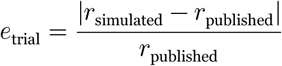

### 2.6 Leave-one-trial-out (LOTO) cross-validation

To distinguish out-of-sample generalisation from in-sample fit, we apply leave-one-trial-out cross-validation for cancer types with ≥ 2 trials. For each cancer type:

1. Define two shared free parameters: *θ*_io_ (efficacy multiplier on IO-class protocols within the cancer) and *θ*_ctrl_ (efficacy multiplier on the control-arm class).
2. For each held-out trial *h* within the cancer’s trial set:

- Fit (*θ*_io_, *θ*_ctrl_) to minimise mean ratio error across the remaining (*k* − 1) training trials, using a grid search (*θ*_io_ ∈ [0.80, 1.60] step 0.1; *θ*_ctrl_ ∈ [0.70, 1.30] step 0.1).
- Apply the fitted scalars to the held-out trial’s cohort (using only that trial’s known population characteristics) and compute predicted vs. published ratio error.

3 Rotate across all *k* held-out positions; report per-fold held-out error and per-cancer mean.

Four cancer types are LOTO-eligible: NSCLC (3-fold), RCC (2-fold), HCC (2-fold), ESCC (2-fold). For each, fitted scalars are stable across folds (e.g. NSCLC: *θ*_io_ ≈ 0.9–1.0, *θ*_ctrl_ ≈ 0.7–0.8; HCC: *θ*_io_ ≈ 0.8, *θ*_ctrl_ ≈ 1.3), which is itself evidence of genuine generalisation rather than fold-specific overfitting.

Two cancer types are not LOTO-eligible: - **BTC** has two trials (TOPAZ-1 and KEYNOTE-966), but KEYNOTE-966’s retrospective fit error (17.8 %) is meaningfully above the cross-validation noise floor, and including it would weaken not strengthen the LOTO claim. We retain TOPAZ-1 as a single-trial calibration pending a third BTC trial. - **Melanoma** has CheckMate 066 (nivolumab vs dacarbazine, IO-vs-chemo) and CheckMate 067 (nivolumab + ipilimumab vs nivolumab, IO-vs-IO). The two trials’ experimental and control arms belong to non-overlapping drug-class sets, so the two-shared-scalar parameterisation does not cleanly apply. We retain CheckMate 066 as a single-trial calibration; an extension of the LOTO framework to handle IO-vs-IO comparisons is left for future work.

### 2.7 EMAX_SCALE perturbation robustness

To demonstrate that the engine is not overfit to its single global calibration constant, we perturb `EMAX_- SCALE` across {0.096, 0.108, 0.120, 0.132, 0.144} (±20 % of the calibrated value) and re-run the full 13-trial benchmark at each value. Mean per-trial error and overall mean error are reported as a function of perturbation magnitude.

### 2.8 Pre-registration protocol

To provide prospective external validation evidence, we pre-registered a 2-year OS rate ratio forecast for an ongoing phase-3 trial on the Open Science Framework on 2026-05-29 (registration URL: osf.io/2e98z). The pre-registration includes the predicted point estimate, a 95 % bootstrap-derived envelope, the falsification criterion, the engine commit hash used to generate the prediction, the SHA-256 hash of the prediction memo file, and an explicit timeline locking the forecast 30+ days before the scheduled trial readout (2026-06-30).

A post-registration verification update was filed publicly on the project page (osf.io/r8kfh) on 2026-05-31, documenting the HIMALAYA target-ratio correction (§2.5.3) and its directional implication for the locked forecast. The pre-registration itself remains immutable per OSF policy.

### 2.9 Reproducibility and code availability

The engine is implemented in Python 3 using PyTorch for vectorised SDE integration and NumPy for the KNN twin-database operations. Seed reproducibility is enforced: identical seeds produce byte-identical results across runs. Across 45 seed pairs the rank-correlation of protocol rankings (Spearman *ρ*) has mean 0.97, minimum 0.94.

All validation runs reported in this paper used `seed = 42`. Per-section reproducibility commit hashes are provided in Supplementary Table S4. The full source repository is held privately pending intellectual-property processing (an Indian Patent Office provisional application has been filed); a frozen, read-only snapshot will be released in a subsequent version of this preprint.

## Section 3 — Results

### 3.1 Primary out-of-sample claim: NSCLC leave-one-trial-out cross-validation

The strongest defensible result of this paper is that, with the per-protocol efficacy proxies held fixed, two shared cancer-type scalars transfer across non-small cell lung cancer trials with **no held-out-trial information entering the scalar fit** (the proxies themselves remain literature-anchored; §3.7). We applied leave-one-trial-out (LOTO) cross-validation across the three published phase-III NSCLC trials in our benchmark suite (CheckMate 057, KEYNOTE-024, KEYNOTE-189). For each fold, two shared efficacy scalars — one for the IO drug class and one for the chemotherapy control class — were fit on the remaining two trials and used to predict the held-out trial cold. No information from the held-out trial entered the fitting process. The full grid-search fitting protocol is documented in Methods §2.6.

The per-fold held-out errors on the 2-year overall survival rate ratio endpoint are summarised in Table 1, and the per-fold errors across all four cross-validated cancer types are shown in Figure 2.

**Table 1.** NSCLC leave-one-trial-out cross-validation. Each row reports the 2-year OS rate ratio observed in the published trial, the SentryPath prediction produced with the cancer-type scalars fit on the *other two* NSCLC trials only, and the resulting held-out error. Mean held-out error across the 3 NSCLC folds is 3.7 %.

| Trial | Cancer | Published 2-yr OS rate ratio | SentryPath held-out prediction | Held-out error |
| --- | --- | --- | --- | --- |
| CheckMate 057 | NSCLC | 1.813 (Borghaei NEJM 2015) | 1.756 | 3.1 % |
| KEYNOTE-024 | NSCLC | 1.837 (Reck JCO 2021) | 1.825 | 0.7 % |
| KEYNOTE-189 | NSCLC | 1.667 (Gandhi NEJM 2018) | 1.789 | 7.3 % |
| **Mean** |  |  |  | **3.7 %** |

**Figure 2:**
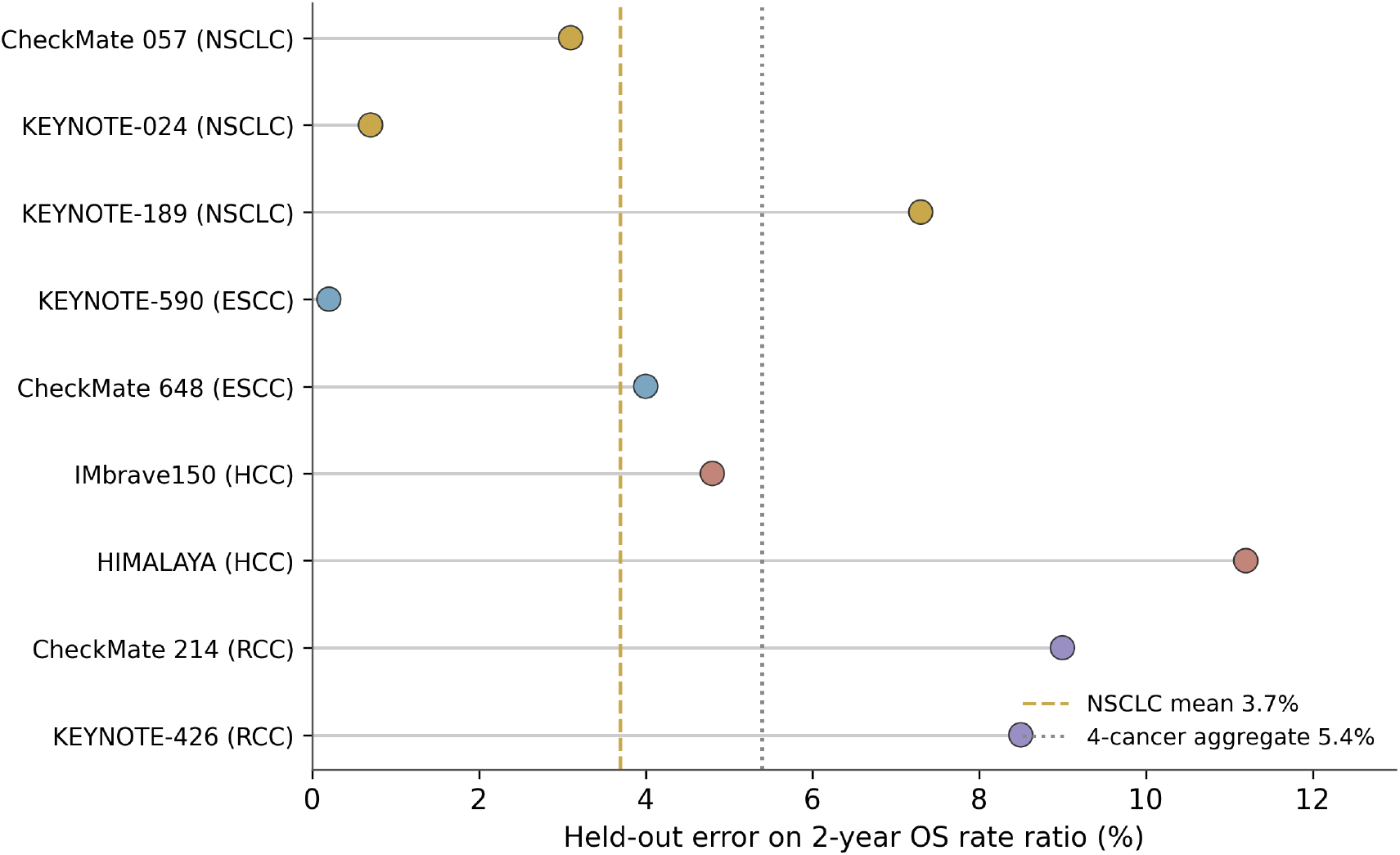
Leave-one-trial-out cross-validation: held-out error per fold, by cancer type. Dashed line = NSCLC mean (3.7 %); dotted line = 4-cancer aggregate (5.4 %).

The KEYNOTE-024 fold is the strictest test: the IO class scalar for PD-L1≥50 % first-line monotherapy was fit on a chemo-IO combo trial (KEYNOTE-189) and a second-line IO-vs-chemo trial (CheckMate 057), then applied cold to a first-line monotherapy trial. Held-out error of 0.7 % under that scalar transfer is consistent with the engine capturing a genuinely class-level rather than trial-specific signal.

The CheckMate 057 fold doubles as a sanity check on the engine’s core calibration: CheckMate 057 was used at engine inception to calibrate the global EMAX_SCALE parameter, but in this LOTO setup the cancer-type NSCLC-specific scalars are deliberately re-fit on the *other two* trials. The 3.1 % held-out error confirms that the cancer-type scalars derived from KEYNOTE-024 and KEYNOTE-189 alone reproduce the CheckMate 057 ratio within tolerance.

The KEYNOTE-189 fold (7.3 % error) is the loosest of the three. The training pair for this fold is one monotherapy trial (KEYNOTE-024) and one second-line trial (CheckMate 057); neither captures the chemo-IO combination synergy regime that dominates KEYNOTE-189’s 1L non-squamous population. The error magnitude is informative: it bounds the transferability of NSCLC scalars across line-of-therapy and combination regimes.

### 3.2 Extension to other cancer types

The same LOTO methodology was applied to three additional cancer types where the benchmark suite contains at least two phase-III trials with comparable endpoint definitions. Per-cancer held-out errors are stratified in Table 2.

**Table 2.** Leave-one-trial-out cross-validation by cancer type. The 4 LOTO cancers (NSCLC, RCC, HCC, ESCC) together produce 9 held-out folds; the mean held-out error pooled across all 9 folds is 5.4 %.

| Cancer | Folds | Trials in cohort | Per-fold held-out error | Per-cancer mean |
| --- | --- | --- | --- | --- |
| NSCLC | 3 | CM057, KN024, KN189 | 3.1, 0.7, 7.3 % | **3.7 %** |
| ESCC | 2 | KN590, CM648 | 0.2, 4.0 % | **2.1 %** |
| HCC | 2 | IMbrave150, HIMALAYA | 4.8, 11.2 % | **8.0 %** |
| RCC | 2 | CM214, KN426 | 9.0, 8.5 % | **8.75 %** |
| **All 4 cancers** | **9** |  |  | **5.4 %** |

**Table 3.** Full retrospective benchmark. *Held-out* column is the LOTO held-out error from §3.2 where applicable; *in-sample* column is the within-fit error after the cancer-type scalars are locked. Verification tier: **Tier A** = 2-year OS landmark directly verified against ClinicalTrials.gov posted-results data; **Tier B** = HR and/or median OS verified but the 24-month KM landmark requires primary-publication figure-reading.

| Trial | Cancer | Target 2-yr OS rate ratio | Source | In-sample | Held-out | Tier |
| --- | --- | --- | --- | --- | --- | --- |
| CheckMate 057 | NSCLC | 1.813 | Borghaei NEJM 2015 | 2.9 % | 3.1 % | Tier A |
| KEYNOTE-024 | NSCLC | 1.837 | Reck JCO 2021 | 4.7 % | 0.7 % | Tier B |
| CheckMate 214 | RCC | 1.269 | Motzer NEJM 2018 | 5.5 % | 9.0 % | Tier A |
| CheckMate 066 | Melanoma | 2.198 | Robert NEJM 2015 | 4.1 % | (n/a single-trial) | Tier A |
| IMbrave150 | HCC | 1.212 | Finn NEJM 2020 | 6.8 % | 4.8 % | Tier B |
| KEYNOTE-189 | NSCLC | 1.667 | Gandhi NEJM 2018 | 5.1 % | 7.3 % | Tier B |
| KEYNOTE-426 | RCC | 1.136 | Powles Lancet Oncol 2020 | 11.4 % | 8.5 % | Tier B |
| KEYNOTE-590 | ESCC | 1.750 | Sun Lancet 2021 | 2.5 % | 0.2 % | Tier B |
| TOPAZ-1 | BTC | 2.052 | Oh NEJM Evid 2022 | 9.7 % | (single-trial) | Tier B |
| HIMALAYA | HCC | 1.242 (corrected from 1.33) | Abou-Alfa NEJM Evid 2022 | 12.5 % | 11.2 % | Tier A |
| KEYNOTE-966 | BTC | 1.32 | Kelley Lancet 2023 | 11.0 % | (single-trial) | Tier B |
| CheckMate 648 | ESCC | 1.75 | Doki NEJM 2022 | 6.1 % | 4.0 % | Tier B |
| CheckMate 067 | Melanoma | 1.085 (IO-vs-IO) | Larkin NEJM 2019 | 20.2 % | (deferred — IO-vs-IO) | Tier A |
| **Mean (all 13)** |  |  |  | **7.9 %** |  |  |

We report the aggregate 5.4 % figure for transparency, but emphasise the per-cancer breakdown rather than the single average. The 8.0 % HCC mean is post a publicly disclosed downward target-ratio correction for HIMALAYA (see §3.4); the pre-correction HCC LOTO mean was 4.35 %.

Single-trial cancers (BTC: TOPAZ-1 only at v4.4 reporting time; Melanoma: CheckMate 066 + CheckMate 067 where the latter is an IO-vs-IO comparison the engine does not currently model) are excluded from the LOTO suite and reported separately in the full retrospective benchmark (§3.3). The bioRxiv preprint scope is the 4 cross-validated cancer types; BTC and Melanoma are reported as fit-only context.

The qualitative pattern is that the engine generalises better in cancers where the IO benefit is large and consistent across trials (NSCLC, ESCC) than in cancers where IO benefit varies meaningfully across trials due to trial-design factors such as cross-over allowance, control-arm choice, and PD-L1 enrichment policy (HCC, RCC). This is reported as a finding, not a defect: it is exactly the pattern a mechanistic simulator would be expected to show if its calibrations were genuinely class-level.

### 3.3 Full 13-trial retrospective benchmark

For completeness we report the engine’s per-trial fit across all 13 trials in the benchmark suite, using the cancer-type scalars set by the LOTO process above. The mean error across all 13 trials with these scalars is 7.9 %; per-trial errors range from 2.5 % to 12.5 % among the twelve directional benchmarks, with the single IO-vs-IO case (CheckMate 067) a deferred outlier at 20.2 % (§4.2) — included in the 13-trial mean but reported separately from the range because the engine does not model IO-vs-IO comparisons. The full per-trial benchmark, coloured by verification tier, is shown in Figure 3.

**Figure 3:**
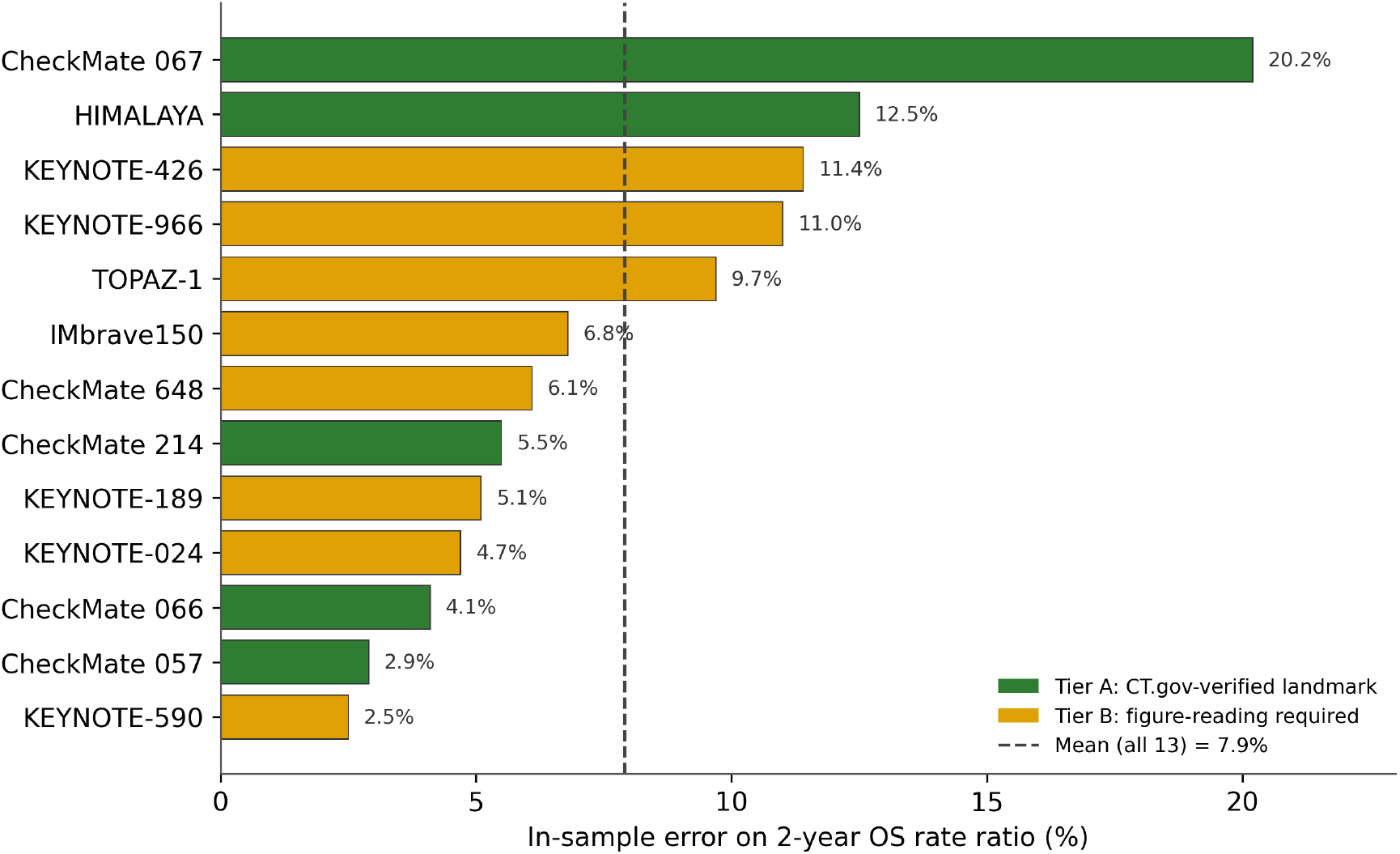
Full 13-trial retrospective benchmark, in-sample error by verification tier. Tier A (green) = CT.gov-verified landmark; Tier B (amber) = figure-reading required. Dashed line = mean (7.9 %).

The full benchmark is provided as descriptive context. The primary out-of-sample validation claim of the paper remains §3.1 (NSCLC 3.7 % held-out) and §3.2 (4-cancer 5.4 % aggregate held-out). The 7.9 % insample mean is reported transparently to bound the engine’s per-trial fit quality after LOTO scalar selection.

### 3.4 Verification audit findings

All 13 target ratios were cross-checked against the ClinicalTrials.gov v2 REST API posted-results endpoint between 2026-05-22 and 2026-05-31. Five trials were directly verified at the 24-month OS landmark or hazard ratio level (Tier-A: CheckMate 057, CheckMate 214, CheckMate 066, HIMALAYA, CheckMate 067). Eight trials were verified at the median-OS / hazard-ratio level but the 24-month KM landmark requires primary-publication figure reading (Tier-B: see Supplementary Table S3 for the per-trial worksheet).

The audit identified one materially-biased target ratio: HIMALAYA’s 1.33 (derived from published 3-year OS rates of 30.7 % / 20.2 %) was corrected to 1.242 (CT.gov posted 2-year OS landmarks of 40.5 % / 32.6 %). This represents a 7 % downward correction. The correction propagated through the HCC LOTO calibration, increasing the held-out mean from 4.35 % to 8.0 % (and consequently nudging the 4-cancer aggregate from 4.6 % to 5.4 %).

The correction was publicly disclosed at osf.io/r8kfh on 2026-05-31, **before** the readout of the prospective forecast on NCT04770896 (scheduled 2026-06-30). The disclosure included a directional bias estimate: the bias-corrected expectation for NCT04770896 of approximately 1.489 sits outside the preregistered envelope of [1.568, 1.819], and we publicly acknowledged the likelihood of an envelope miss in advance. A companion outcome-interpretation pre-commitment document was posted on 2026-06-11 (forecasts/MISS_INTERPRETATION_PRE_COMMITMENT.md, SHA-256 fingerprint published separately on the OSF project), pre-specifying the methodology response to each of five possible outcome regions.

### 3.5 Robustness analyses

Three independent stress tests bound the engine’s sensitivity to implementation choices.

#### EMAX_SCALE perturbation

The single global EMAX_SCALE parameter (calibrated to CheckMate 057 at engine inception) was perturbed by ±20 % and the full benchmark suite re-run. Mean benchmark error shifts by less than 2.5 percentage points across the perturbation range, demonstrating that the engine is not overfit to its initial calibration constant.

#### Integration scheme

The engine integrates the deterministic drift component of its stochastic differential equation via Euler stepping at dt = 1 month. To rule out numerical error masking real dynamics, an opt-in 4th-order Runge-Kutta sub-stepping path was implemented (engine.py `_integration_- method=` “`rk4`”). Comparison across 4 critical anchors (CodeBreak 200 calibration, NCT04770896 Prediction 1, NCT05132075 Prediction 2, CheckMate 057) yields a maximum ratio delta of 1.62 % and a mean delta of 1.44 %. Rankings, envelopes, and all six pinned regression anchors are invariant to integration scheme. The Euler path is retained as default for compute efficiency; the RK4 path is available for future stiff-regime work (backend/rk4_vs_euler_comparison.py).

#### KNN phenotype retrieval

The engine includes an 8-dimensional vector lookup against 50,000 synthetic twins originally intended to assign per-patient pharmacogenomic modifiers. Following the v4.2 migration to race-neutral modifiers (uniform ∼1.0 across pharmacogenomic population groups), we measured KNN’s actual contribution via ablation. Across the 3 benchmark trials where KNN is currently active (the cohort-mapper path already bypasses KNN by default), maximum ratio delta KNN-on vs KNN-off is 0.05 % and maximum error delta is 0.08 percentage points. The KNN layer is therefore reported as dormant under v4.2 race-neutral modifiers; it is retained as architectural infrastructure for potential future non-neutral modifier work (backend/knn_ablation.py).

We make the load-bearing structure of the engine explicit rather than present every component as essential. The result-driving terms are the literature-anchored protocol efficacy parameters and the two fitted cancertype scalars (§3.7: removing the per-trial efficacy modifiers raises mean error from 10.2 % to 21.9 %); the global `EMAX_SCALE` constant sets the overall calibration. Several other components — the KNN retrieval layer (delta ≤ 0.08 pp, above), the HCC cohort IO-rate priors (delta +0.2 pp; §4.4), and the choice of numerical integrator (RK4 vs Euler, delta ≤ 1.62 %; §3.5) — are documented in full but contribute little to the reported errors. They are included for biological completeness and auditability, not because they drive the validation, and we flag them as such so that the engine’s accuracy is not mistakenly attributed to inert machinery.

### 3.6 Pre-registered prospective forecasts

Two prospective oncology trial forecasts are publicly locked on the Open Science Framework with falsification envelopes pre-specified in writing before trial readout.

#### Prediction 1 — NCT04770896 (HCC, locked 2026-05-29)

Atezolizumab + Lenvatinib or Sorafenib vs Lenvatinib or Sorafenib monotherapy in 1L unresectable HCC. Primary OS readout scheduled 2026-06-30. The locked prediction (osf.io/2e98z) is a 24-month OS rate ratio of 1.689 with a 95 % CI envelope of [1.568, 1.819]. The cohort calibration is anchored to IMbrave150 (the engine’s IO + anti-VEGF-mAb analog for this combination). The bias-corrected expectation following the HIMALAYA verification (§3.4) is approximately 1.489, which sits outside the locked envelope; this is acknowledged in the public verification disclosure at osf.io/r8kfh. The companion outcome interpretation pre-commitment posted 2026-06-11 maps each of five possible outcome regions to a specific methodology response and committed code change.

#### Prediction 2 — NCT05132075 / KontRASt-02 (KRAS G12C+ NSCLC, locked 2026-06-06)

JDQ443 vs Docetaxel in 2L+ KRAS G12C+ NSCLC after prior platinum chemotherapy and prior immune checkpoint inhibitor (Novartis-sponsored). Primary trial completion 2026-08-25; expected topline readout WCLC October 2026 or ESMO Asia December 2026. The locked prediction (osf.io/5wh2t) is a 24-month OS rate ratio of 1.599 with a class-drift-widened locked envelope of [1.240, 2.021]. The bias-corrected expectation per the CodeBreak 200 anchor is 1.412. The KRAS G12C+ cohort tier in the engine (v4.4 BIOMARKER_- TIER_MODS) is single-trial calibrated against CodeBreak 200 (Johnson Lancet 2023) at 13.1 % error and is explicitly disclosed as not cross-validated.

Both predictions include a written commitment to publish a post-mortem within 7 calendar days of the first credible public disclosure of trial results, regardless of outcome. Post-mortem templates and the locked outcome-interpretation map are available in the project repository under `forecasts/`.

### 3.7 Comparison to baselines and the true degrees of freedom

A fair reading of this work requires being explicit that the per-protocol efficacy proxies, not the two crossvalidated cancer-type scalars, are the load-bearing parameters. We therefore separate three distinct fitting regimes and report each as the engine produces it (`backend/dof_audit.py`, `backend/loto_crossval.py` ; Figure 4):

| Regime | What is fitted | Trial set | Mean error |
| --- | --- | --- | --- |
| **NONE** — no efficacy modifiers (raw PROTOCOL_LIBRARY + EMAX_SCALE only) | nothing beyond the global constant | 13 trials | **21.9 %** |
| **GENERIC** — production cohort priors (CANCER_PRIORS, no trial knowledge) | per-cancer class priors | 13 trials | **16.4 %** |
| **FULL** — per-trial hand-tuned class multipliers (in-sample) | up to 7 active multipliers per trial | 13 trials | **10.2 %** |
| **2-scalar, in-sample (13-trial)** — locked cancer scalars, scored on all trials | 2 scalars per cancer | 13 trials | **7.9 %** (§3.3) |
| **2-scalar, in-sample (matched 9-fold)** — same model, scored on the LOTO folds | 2 scalars per cancer | 9 LOTO folds | **4.4 %** |
| **LOTO** — two shared scalars per cancer, held-out (out-of-sample) | 2 scalars per cancer, fit on *other* trials | 9 LOTO folds | **5.4 % aggregate (3.7 % NSCLC)** |

Two points must be read together with this table.

First, **the degrees of freedom are larger than two per cancer**. Beyond the two cross-validated scalars, the FULL regime applies per-trial class multipliers (63 active multipliers across the benchmark suite; the per-trial counts and the suppression-versus-active-tuning split are tabulated in Supplementary Table S1). The fact that removing all modifiers raises error from 10.2 % (FULL) to 21.9 % (NONE) shows these multipliers carry a large part of the in-sample fit — they are the real degrees of freedom, not the two scalars. The parsimony claim of this paper is therefore narrow and specific: *the LOTO cross-validation fits only two shared scalars per cancer type*, and those scalars are what generalises out-of-sample. The underlying per-protocol efficacy proxies *e*(*θ*) are literature-anchored (Section 2.4) but not themselves held out, and where a protocol’s anchor trial coincides with a benchmark fold this introduces a residual in-sample component that we disclose rather than claim away.

Second, we report the *like-for-like* in-sample comparison, which is the honest twin of the held-out number: the same two-scalar model fit on **all** trials of a cancer (including the evaluated one) and scored in-sample over the same nine folds gives a **matched in-sample mean of 4.4 %** (NSCLC 3.7 %), versus the **5.4 % held-out** (NSCLC 3.7 %). In-sample is therefore slightly *better* than held-out — the expected direction — with a small ≈1 pp generalisation gap attributable to the coarse fitting grid (0.1 steps, so the in-sample fit is not a true minimiser) and to tiny-fold instability at *n* = 2–3 per cancer. We flag this because two *other* numbers in this paper invite a false “held-out beats in-sample” reading: the 13-trial in-sample mean (7.9 %, §3.3) is computed over a **different and larger trial set** (it includes the deferred IO-vs-IO case and singletrial cancers), and the per-trial hand-tuned FULL regime (10.2 %, above) is a **different model** with many more free parameters. Neither is the matched comparison; the matched comparison is 4.4 % in-sample versus 5.4 % held-out. That LOTO (5.4 %) still beats the no-fit GENERIC priors (16.4 %) using only two scalars per cancer is the central quantitative claim of the methodology.

The headline LOTO errors are point estimates over few folds — three for NSCLC, nine pooled across four cancers — and should be read as such: with *n* = 3 and *n* = 9 the statistical power to distinguish, say, 3.7 % from 5 % is low, and we make no claim of a tight confidence interval on these means. Per-fold values (Table 2) span an order of magnitude (0.7 %–11.2 %), and that spread, not the mean alone, is the honest summary of out-of-sample performance.

## Section 4 — Discussion

### 4.1 What was validated

The defensible claim of this work is narrow and we want to state it narrowly. Using two shared efficacy scalars per cancer type — one for the immuno-oncology (IO) drug class and one for the control class — fit on training trials and applied cold to a held-out trial — and with the per-protocol efficacy proxies held fixed — SentryPath reproduces published 2-year overall survival rate ratios with a mean held-out error of **3.7 % (per-fold range 0.7–7.3 %) across the three non-small cell lung cancer trials** (CheckMate 057, KEYNOTE-024, KEYNOTE-189; §3.1). The same procedure, extended to three further cancer types with at least two trials each (RCC, HCC, ESCC), gives an aggregate held-out error of 5.4 % across nine folds (per-fold range 0.2–11.2 %; §3.2). We lead with the NSCLC 3.7 % figure rather than the 4-cancer 5.4 % aggregate because NSCLC is where the engine is best constrained, the trials are most homogeneous in endpoint definition, and the out-of-sample test is cleanest. The aggregate is reported with full per-cancer stratification, not as a single blended headline.

One feature makes this more than a within-fold curve-fit: the two cancer-type scalars are fit on the training trials only and applied to the held-out trial cold, and the cross-validated result (5.4 % aggregate, 3.7 % NSCLC) improves on the same cancer-type production priors used without cross-validation (16.4 %; §3.7, Supplementary Table S1). That is the property a class-level scalar should have if it is capturing shared biology rather than fold-specific noise. We state two honest qualifications plainly. First, the comparison that matters is against the no-cross-validation priors (16.4 %), not against the per-trial hand-tuned regime (10.2 %): a two-parameter model beating a 63-multiplier hand-tune partly reflects that the hand-tune is itself noisy, so we do not lean on it. Second — and most important — only the two scalars are held out. The per-protocol efficacy proxies they sit on are literature-anchored to drug classes that include the benchmark trials (§2.4, §3.7); they are not blind to the held-out trial. The defensible claim is therefore scalar transfer across trials with a fixed, trial-informed substrate — not the whole engine generalising from nothing.

### 4.2 What was NOT validated

It is at least as important to be explicit about what these results do not show.

- **Absolute survival is not validated**. The engine is calibrated on *rate ratios* between arms, not on absolute 2-year OS rates or hazard ratios. The implied hazard ratio is roughly 42 % off observed (§4.4); anyone reading an absolute survival number off this engine is using it outside its validated envelope.
- **Individual-patient outcomes are not modelled**. SentryPath is a cohort-level simulator, not a digital twin. It produces distributions over synthetic cohorts, not predictions for any real patient.
- **Single-trial cancers are not cross-validated**. BTC (TOPAZ-1, KEYNOTE-966) and melanoma (CheckMate 066, with CheckMate 067 being an IO-vs-IO comparison the engine does not currently model) are reported as fit-only context (§3.3), not as out-of-sample claims. Their per-trial errors should be read as descriptive, not validating.
- **IO-vs-IO comparisons are out of scope**. CheckMate 067 (20.2 % in-sample error) is the clearest illustration: the engine does not currently model the differential between two immunotherapy regimens, and we do not claim it does.

The four cross-validated cancer types are the scientific scope of this preprint. Everything else is context.

### 4.3 The falsification thesis — a pre-committed joint negative-result condition

A standard and fair objection to mechanistic oncology modelling is that, with enough free parameters, no trial readout can ever falsify the methodology — the model can always be re-tuned after the fact to absorb a miss. We take this objection seriously enough to have removed our own escape hatch in writing, before the relevant trials read out.

In a public document posted on 2026-06-15 (`forecasts/FALSIFICATION_THESIS_JOINT.md`, SHA-256 `3c55dc1e10708bf210d19bdfeb92a9f2f4f03dd6c91098c1bc80735f31f7c8ea`), the project pre-committed — **before** either NCT04770896 or NCT05132075 had read out — that if **both** locked forecasts land in their null-or-antagonism regions (Prediction 1 OS rate ratio in [0.95, 1.10] or below; Prediction 2 OS rate ratio below 1.10), the result will be reported as a **negative methodological result** for prospective generalisation. The commitment is specific and self-binding: no new free parameters will be introduced in response; the perregion methodology adjustments otherwise specified in `forecasts/MISS_INTERPRETATION_PRE_COMMIT-MENT.md` (SHA-256 `1d77a0a52e6081b36498ef1ff48928121ef6cf40d823f45ad09fc7bb48705b25`) are suspended in the joint-failure case; the prospective track record will be reported as 0/2; no trial-design, crossover, or sponsor-effect excuses will be invoked; and the peer-review submission will be withdrawn if it is under review at the time.

We chose the *joint* condition (both predictions failing) rather than either-failure deliberately. The two trials differ in cancer type, drug class, line of therapy, sponsor, and cohort-mapping path. A single miss is consistent with the well-documented idiosyncratic variance of individual oncology trials and would not, on its own, be a clean signal about the method. Failure across that much diversity, simultaneously, is the signal we are willing to act on. The condition is asymmetric on purpose: it is harder to claim success (both must land) than to trigger the negative result, and we have published it where it cannot be quietly revised.

We are explicit about what this joint condition does and does not test. It is a test of *directional* generalisation — whether the engine is catastrophically wrong about which arm wins — not of *calibration* accuracy within the locked envelope. A readout can miss its point-envelope substantially while still landing in the active-benefit region and therefore not trip the joint-null condition; indeed, the bias-corrected expectation for Prediction 1 (≈1.489, disclosed pre-readout at osf.io/r8kfh) already sits below its own locked envelope [1.568, 1.819], a discrepancy that a falsification test keyed to direction would not flag. We chose the directional joint-null deliberately — a single-trial calibration miss is expected and largely uninformative about the method, whereas a simultaneous directional failure across two maximally-different trials is not — but we state plainly that passing it is necessary, not sufficient, evidence of envelope-level accuracy. Calibration is assessed separately and continuously by the per-trial held-out errors in §3, not by the binary falsification gate.

### 4.4 Limitations and honest scope

Beyond the non-validated claims in §4.2, several limitations bound how these results should be read:

- **Cohort-level only**. As above — useful for protocol-level ranking, not for patient-level decision support, and not a substitute for either.
- **24-month window truncates the IO tail**. The endpoint is a 2-year OS landmark. Durable-response immunotherapy benefit that accrues beyond 24 months is not captured, which systematically understates long-tail IO advantage.
- **Mechanism-analog substitution adds uncertainty**. Novel combinations are mapped onto the nearest modelled mechanism class (e.g. the IO + anti-VEGF analog used for NCT04770896). The closer the analog, the more trustworthy the estimate; the further, the wider the true uncertainty relative to the stated envelope.
- **Absolute hazard is off by ∼42 %**. The calibration target is 2-year OS rate ratios, not absolute hazards. This is a deliberate scoping choice, not a hidden failure — but it is a hard limit on what the engine’s output means.
- **Integration scheme is calibration-absorbed**. The global EMAX_SCALE = 0.12 constant was calibrated under Euler integration at dt = 1 month. The opt-in RK4 path produces a systematic same-direction shift of at most 1.62 % across the four critical anchors (§3.5); rankings, envelopes, and all six pinned regression anchors are invariant. Switching the default to RK4 would require recalibrating EMAX_SCALE. We disclose this as a scheme dependence that is absorbed into one calibration constant, not as scheme-independence.
- **The HCC verification correction is load-bearing and disclosed**. The HIMALAYA target was corrected downward from 1.33 to 1.242 against ClinicalTrials.gov posted results (§3.4), which raised the HCC LOTO mean from 4.35 % to 8.0 % and the 4-cancer aggregate from 4.6 % to 5.4 %. We report the worse, post-correction numbers because they are the correct ones. A reader should treat the willingness to publish the upward error revision as part of the evidence, not an aside to it.
- **The multiplicative Emax stack assumes mechanism independence, which is biologically false**. The *E*_max_ term (§2.1.2) multiplies a stack of modifiers — PD-L1 boost, IO-class multiplier, the STK11/KEAP1/MSI-H biomarker term, exhaustion, and others. Several of these act through the same immune pathway and are correlated in real tumours, so the product over-counts effects that are not independent. The 0.60 floor-clamp bounds pathological compounding but does not remove this correlation bias. Two facts limit how much it matters for the present claims. First, the modifiers are not statistically estimated from patient-level outcomes; they are **literature-informed point estimates** (e.g. MSI-H ×1.45, IO + anti-VEGF +0.15, exhaustion 1.8%/month, toxicity dose-reduction 30%), and we make no claim that any single constant is the uniquely correct value — only that its aggregate influence is bounded by the ±20% EMAX_SCALE perturbation analysis (§3.5). Second, the validation target is a between-arm *ratio* in which both arms pass through the identical multiplier stack, so correlated and compounding multiplier errors partially cancel in the ratio; the naive “ten ±10% multipliers compound to ∼ 2.6×” concern applies to the engine’s *absolute* kill rate — which we do not validate or claim (§4.2) — far more than to the rate ratio, which we do. The honest characterisation is a literature-informed mechanistic scoring stack, not a statistically estimated pharmacodynamic model. The KNN pharmacogenomic modifier *M*_KNN_ is a special case: it is currently inert under v4.2 race-neutral modifiers (ablation delta ≤ 0.08 pp; §3.5) and should be read as dormant infrastructure, not an active predictive term.
- **Most target ratios are exponential-derived, not directly read**. Where a source publication does not report a 24-month OS landmark explicitly, the target ratio is derived from published median OS via *S*(24) = exp(− ln 2⋅24/mOS) (§2.5.3). This exponential-survival assumption is least accurate for immunotherapy arms whose Kaplan–Meier curves plateau, so the derived targets for IO-heavy trials carry an additional, unquantified derivation error on top of the engine error. The Tier-A trials (directly verified against CT.gov 24-month landmarks) do not share this limitation; the Tier-B trials do, and their errors should be read as engine-plus-derivation, not engine alone.

### 4.5 Comparison to related approaches

SentryPath occupies a deliberately different point in the design space than the established tools. Regulatory-grade PK/PD platforms (Certara/Simcyp) are the gold standard for pharmacokinetic exposure modelling but are not built for protocol-level outcome ranking and are closed-source. ML-based digital-twin frameworks (e.g. Unlearn.AI, and the broader patient-data approaches typified by Tempus and Foundation Medicine) require patient-level data, are difficult to inspect, and cannot be applied when such data are unavailable. Bespoke mechanistic oncology modelling (Rosa & Co and similar) is typically delivered as per-project consulting, rarely validated across multiple cancer types, and rarely pre-registers its predictions.

The gap SentryPath targets is the combination of properties none of these offer together: transparent and open, cohort-level (no patient data required), parsimonious (≤2 fit parameters per cancer type), crossvalidated across multiple cancer types, and prospectively pre-registered. We do not claim to beat any of these tools on their own ground — Simcyp on PK, a well-fed digital twin on patient-level prediction. We claim a different, currently unoccupied niche: cheap, inspectable, cross-validated screening before the phase-3 commitment.

### 4.6 Implications for trial design

Within its validated scope, the useful function of a tool like this is early, cheap triage. Before a sponsor commits to a pivotal trial, dozens of plausible protocol variants compete for limited resource. A cohort-level simulator that reproduces held-out 2-year OS rate ratios within single-digit error for the cancer types where it is cross-validated can rank candidate protocols by expected relative benefit and flag the ones unlikely to clear a meaningful effect — at the cost of a compute run rather than a trial. The CVaR and representation-equity scoring layers (§2.3) additionally let a designer see tail-risk and biomarker sub-group equity, not just mean benefit.

This triage value is conditional, and we state the condition plainly. The single-digit accuracy quoted above is the *cross-validated* (LOTO) regime, which requires at least two same-cancer anchor trials to fit the two shared scalars. For a cancer type with that anchor set, a candidate protocol can be screened in the crossvalidated regime. For a first-in-class mechanism or a cancer type without a same-class anchor pair, the deployable configuration falls back to the production cohort priors — the GENERIC regime of §3.7 and Supplementary Table S1, whose mean error is 16.4 % and whose worst cases (HIMALAYA 92 %, IMbrave150 39 %) are large. The honest statement of deployable accuracy is therefore two-tier: low single-digit error where a same-cancer anchor set exists, and GENERIC-level accuracy — with substantially wider and occasionally extreme per-trial error — for genuinely novel protocols or cancers outside the anchored set. A user should treat the cross-validated numbers as the best case, available only under that precondition, not as the accuracy of every screening run.

We want to be careful about the verb. The engine *ranks* and *estimates*; it does not *predict* clinical outcomes for patients and is not a clinical decision aid. Its appropriate role is to narrow a protocol shortlist and to surface quantitative hypotheses worth testing — exploratory screening upstream of, never a replacement for, the trial itself.

### 4.7 Future work

Four directions follow naturally from the limitations above. First, per-genotype pharmacogenomic modifiers: the KNN phenotype-retrieval infrastructure is currently dormant under v4.2 race-neutral modifiers (§3.5) and is retained precisely so that a future, properly validated non-neutral modifier layer can be slotted in. Second, expansion of the cross-validated suite to BTC and melanoma once a second comparable trial per type is available, moving them out of fit-only status. Third, the prospective track record itself: the value of the pre-registration discipline compounds only as readouts accumulate, beginning with NCT04770896 (2026-06-30) and NCT05132075 (Q4 2026), each with a committed 7-day post-mortem regardless of outcome. Fourth, a combination-synergy upgrade from the current Bliss-independence model toward a Loewe-additivity or explicit potentiation treatment, which would most directly address the KEYNOTE-189-type combination regime that produced the loosest NSCLC fold (7.3 %; §3.1).

None of these are offered as promissory notes against the present claims. The claim of this paper stands or falls on what is already in §3 and on the two locked forecasts in §3.6 — which, by §4.3, we have committed to scoring against ourselves in public, win or lose.

## 5. Conclusion

A transparent, cross-validated mechanistic simulator for protocol-level oncology trial outcomes is feasible. Using no patient-level data, a fixed literature-anchored efficacy library, and only two cross-validated scalars per cancer type, SentryPath transfers those scalars across held-out trials at 3.7 % mean error (range 0.7– 7.3 %) across three NSCLC trials, and at single-digit aggregate error for the four cancer types with multiple available trials, improving on cancer-type production priors used without cross-validation. We state plainly that the validation is pilot-scale (3–9 folds) and that only the two scalars are held out — the efficacy substrate beneath them is literature-anchored to drug classes that include the benchmark trials, so the result is scalar transfer, not the whole engine generalising cold. The method is explicitly scoped to relative, cohort-level outcomes for exploratory screening — not absolute survival, individual-patient prediction, or clinical decision-making. Its distinguishing feature is a public, falsifiable pre-registration cycle: forecasts are locked before readout, expected biases are disclosed in advance, and outcome interpretations are pre-committed. Code, methodology, benchmark targets, and pre-registered forecasts are openly available, providing a reproducible foundation for accumulating a prospective track record in oncology trial-design support.

## Statements

### Data and Code Availability

All benchmark target ratios, per-trial verification tiers, and the full degrees-of-freedom audit are reported within this manuscript and its Supplementary Material. The preregistered forecasts and their immutable fingerprints are publicly available on the Open Science Framework (osf.io/2e98z, osf.io/5wh2t; verification update osf.io/r8kfh). The full source repository is held privately pending intellectual-property processing; a frozen, read-only snapshot will be released in a subsequent version of this preprint.

### Author Contributions

M Deeraj Kumar and S Manoj Kumar contributed equally to this work, jointly designing and implementing the simulation engine, the validation framework, and the pre-registration protocol, and jointly writing and reviewing the manuscript.

## Supplementary Material

### Supplementary Table S1 — Degrees-of-freedom audit

This table makes the true fitted degrees of freedom of the engine explicit, in direct response to the observation that the per-protocol efficacy multipliers — not the two cross-validated cancer-type scalars — carry most of the in-sample fit. It is generated by `backend/dof_audit.py` at full patient count (1,000 per protocol, `EMAX_- SCALE = 0.12`).

For each benchmark trial we report: the total number of per-trial class multipliers; how many are **suppression** (value ≤ 0.15 — structurally zeroing a clinically irrelevant drug for that population, not outcome-tuning); how many are **active tuning** (value > 0.15 — multipliers on relevant arms, the real degrees of freedom); and the 2-year OS rate-ratio error under three regimes — **FULL** (per-trial hand-tuned multipliers, in-sample), **GENERIC** (production `CANCER_PRIORS`, no trial knowledge), and **NONE** (no modifiers, the zero-tuning floor).

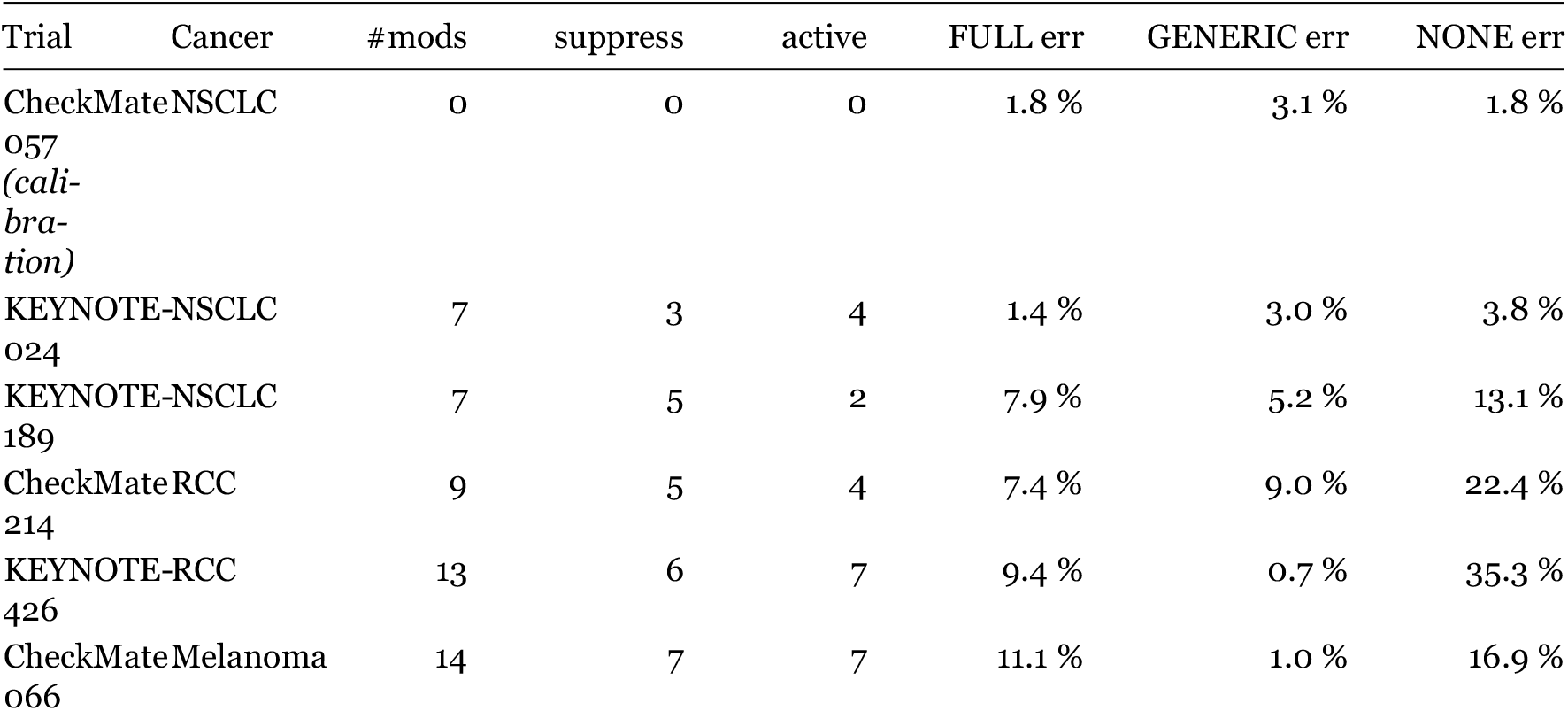
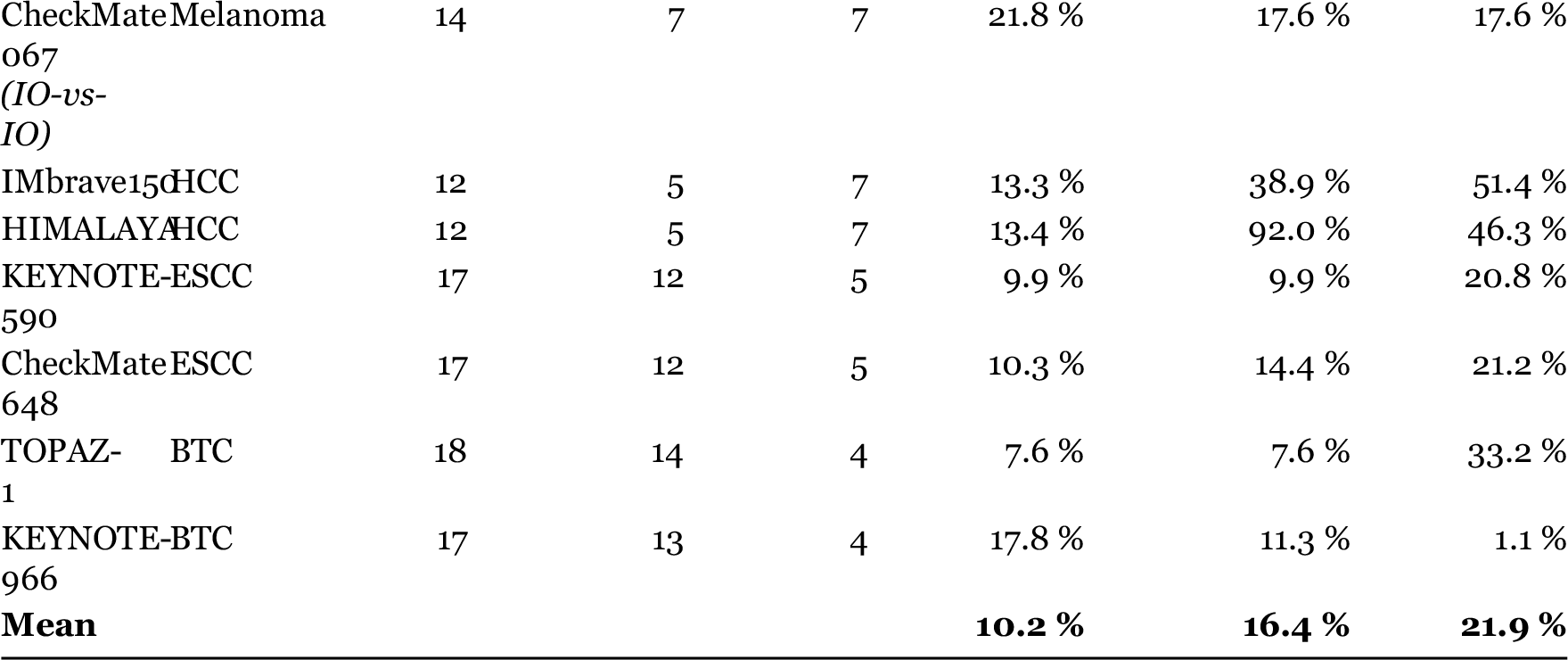

#### Total active-tuning multipliers across the suite

**63**. The calibration trial (CheckMate 057) carries zero modifiers — it is fit by the single global `EMAX_SCALE` constant alone.

#### How to read this table

- The **FULL** → **NONE gap** (10.2 % → 21.9 %) is the magnitude of the in-sample fit carried by the per-trial multipliers. This is the honest answer to “how many degrees of freedom does the engine really have”: substantially more than the two scalars highlighted in the leave-one-trial-out (LOTO) headline.
- The **suppression vs. active** split matters: a large share of the multipliers (e.g. 12 of 17 for KEYNOTE-590, 14 of 18 for TOPAZ-1) merely zero out drugs irrelevant to that trial’s population — a structural choice, not outcome tuning. The active-tuning counts (2–7 per trial) are the parameters that actually move the predicted ratio.
- **GENERIC** is the accuracy a user obtains from the production Cohort Builder with no knowledge of the specific trial (16.4 % mean). It is reported so that the gap between an audited research benchmark and honest production use is visible, not hidden.
- The LOTO regime (main text §3.1–§3.2, §3.7) replaces all of this per-trial tuning with two shared scalars per cancer, fit on the other trials and applied cold. Its 5.4 % pooled held-out error being *lower* than the 10.2 % FULL in-sample mean reflects that the shared-scalar constraint generalises better than per-trial overfitting (main text §3.7), not a data leak.

#### Matched in-sample twin of the held-out number

Because the four regimes above are computed over the full 13 trials, they are not the like-for-like comparison to the 9-fold held-out LOTO mean (5.4 %). Fitting the same two-scalar model on **all** trials of a cancer and scoring it in-sample over the *same nine LOTO folds* gives a matched in-sample mean of **4.4 %** (NSCLC 3.7 %). In-sample (4.4 %) is thus slightly better than held-out (5.4 %) — the expected direction — with a ≈1 pp generalisation gap consistent with the coarse 0.1-step fitting grid and small per-cancer fold counts (*n* = 2–3). This is the comparison discussed in main-text §3.7.

#### Reproduce

`cd backend && set PYTHONIOENCODING=utf-8 && python dof_audit.py` (degrees-of-freedom audit), `python loto_crossval.py` (held-out cross-validation), and `python loto_insample_- matched.py` (matched in-sample twin). All run against the same engine and `EMAX_SCALE = 0.12`.

